# *Neisseria gonorrhoeae* diagnostic escape from a *gyrA*-based test for ciprofloxacin susceptibility can lead to increased zoliflodacin resistance

**DOI:** 10.1101/2022.08.10.503517

**Authors:** Daniel H. F. Rubin, Tatum D. Mortimer, Yonatan H. Grad

## Abstract

The etiologic agent of gonorrhea, *Neisseria gonorrhoeae*, has become resistant to each of the first line antibiotics used to treat it. However, most isolates in the US remain susceptible to ciprofloxacin, leading to the hypothesis that a diagnostic that accurately reports susceptibility to ciprofloxacin may allow its reintroduction into clinical use. One diagnostic approach is to identify codon 91 in *gyrA*, where coding for the wildtype serine is associated with ciprofloxacin susceptibility and phenylalanine with resistance. Here, we found that escape from this diagnostic method can occur through *gyrA* F91S reversion and expansion of pre-existing ciprofloxacin non-susceptible *N. gonorrhoeae* lineages with *gyrA* 91S. Furthermore, we found that mutations conferring ciprofloxacin resistance in *gyrA* 91S isolates led to cross-resistance with zoliflodacin, an antibiotic in phase 3 trial for treatment of gonorrhea. These findings define *N. gonorrhoeae* mutations that should be monitored via surveillance systems, particularly in populations and regions deploying *gyrA* 91-based diagnostics, and demonstrate how diagnostics that guide antibiotic therapy can have unintended consequences.

## Introduction

*Neisseria gonorrhoeae*, the causative agent of the sexually transmitted disease gonorrhea, has developed resistance to each of the first-line antibiotics used to treat it^1^. This rise in antibiotic resistance has led to fears of untreatable highly drug resistant gonorrhea^2^, a concern further heightened by the international circulation of strains resistant to the current first-line antibiotic, ceftriaxone^3–5^.

Due to high levels of drug resistance in *N. gonorrhoeae*, there has been substantial interest in the development of new approaches for management of gonorrhea^6^. One promising avenue has been the use of point-of-care diagnostics that test for susceptibility. By understanding alleles associated with antibiotic susceptibility^7^, sequence-based diagnostics may enable diagnosis of gonorrhea and inform treatment decisions by using genotype to predict antibiotic susceptibility phenotype.

An early target for this approach has been the fluoroquinolone class antibiotic ciprofloxacin^8,9^. Once first-line treatment of *N. gonorrhoeae* infections, rising levels of resistance and concern for failure of empiric therapy resulted in recommendations against its empiric use^10^. However, surveillance data from the US, the UK, and Europe indicate that ciprofloxacin resistance is present in a minority of cases^11–13^, suggesting that a diagnostic that can inform on ciprofloxacin susceptibility would allow for the drug’s reintroduction to gonorrhea treatment and thereby aid in reducing the selective pressure on ceftriaxone^14^.

Ciprofloxacin resistance in *N. gonorrhoeae* is mainly mediated by mutations in the A subunit of DNA gyrase (encoded by *gyrA*) and the A subunit of topoisomerase IV (encoded by *parC*). Though multiple mutations contribute to ciprofloxacin resistance, including position 95 in GyrA and multiple sites at positions 86-91 in ParC^15^, position 91 in GyrA is highly predictive of clinical susceptibility (if a serine) and resistance (if a phenylalanine) to ciprofloxacin^16,17^.

The SpeeDx ResistancePlus GC diagnostic test reports the presence of *N. gonorrhoeae* and ciprofloxacin susceptibility. The ResistancePlus GC test consists of a multiplex qPCR, incorporating channels for the wild-type (WT) *gyrA* 91S allele, the mutant *gyrA* 91F allele, the *N. gonorrhoeae opa* gene, the *N. gonorrhoeae porA* gene, and a PCR positive control. Across multiple studies, the ResistancePlus assay detects gonorrhea with a sensitivity of 94-100%^8,18,19^ and specificity of 99-100%^8,19^. The test is also highly sensitive (>99%) and specific (>97%) for ciprofloxacin resistance^18^.The ResistancePlus GC assay has been approved in Europe and received a breakthrough designation from the FDA^20^.

The use of sequence-based diagnostics raises the possibility of diagnostic escape, with a precedent in *N. gonorrhoeae* of mutations that resulted in failure of clinical identification by PCR ^21,22^. In this case, the diagnostic applies selective pressure for *N. gonorrhoeae* to develop resistance through a pathway other than the *gyrA* 91 mutation it targets. False negative tests for antibiotic susceptibility testing may result in clinical treatment failure for the individual and increased burden of disease and accelerated spread of resistance in the population^23^. The risk of ciprofloxacin resistance arising through novel pathways is especially pressing due to concern for cross-resistance to the new *N. gonorrhoeae* antibiotic zoliflodacin, a first-in-class spiropyrimidinetrione antibiotic that primarily targets the B subunit of DNA gyrase (GyrB)^24^ and is in Phase 3 trials for treatment of gonorrhea^25^.

In this study, we investigated the pathways of diagnostic escape from tests that rely on GyrA 91. We found that some clinical isolates with a 91S allele of GyrA maintain ciprofloxacin resistance at a level that has been associated with treatment failure, and that one clinical isolate (GCGS0481) in which we replaced GyrA 91F with 91S acquired high-level ciprofloxacin resistance while maintaining the 91S allele. Finally, though these strains are zoliflodacin naïve, we found that, in the presence of ciprofloxacin, clinical isolates readily evolve a D429N mutation in GyrB that confers ciprofloxacin resistance and that has been reported to confer zoliflodacin resistance^26^. Taken together, these results indicate that there are evolutionary pathways to diagnostic escape from *gyrA* 91-based diagnostics and that one of these pathways leads to cross-resistance between ciprofloxacin and zoliflodacin. Finally, these results provide experimental validation for the hypothesis that single antibiotic sequence-based diagnostics may lead to unanticipated consequences, including novel resistance determinants and cross-resistance.

## Methods

### Bacterial strains, MIC measurements and Cloning

The strains used in this study are listed in Supplementary Table 1. *N. gonorrhoeae* was grown on GCB Agar (Difco) supplemented with Kellogg’s supplements^27^ at 37°C in 5% CO2. Antibiotic susceptibility testing was performed on GCB-K agar via agar dilution for zoliflodacin (using the range 0.008 to 4 μg/mL) and Etest (BioMerieux) for ciprofloxacin, ceftriaxone, erythromycin, benzylpenicillin, and tetracycline.

Cloning was performed using primers and plasmids listed in Supp. Table 1 and Gibson assembly ^28^ into a pUC19^29^ backbone using a kanamycin cassette from pDR1^30^. Fragments were amplified using Phusion (NEB), checked for appropriate size by gel electrophoresis, column purified (Qiagen PCR Purification kit), assembled with Gibson Master Mix (NEB), and finally transformed into chemically competent DH5α *E. coli*. Finally, individual colonies were selected on LB agar with 50 μg/mL kanamycin, picked, and grown overnight. Plasmids were isolated using spin columns (Qiagen Spin Miniprep Kit). The resulting plasmids were checked by Sanger sequencing. For insertion of GyrA alleles into *N. gonorrhoeae*, strains of *N. gonorrhoeae* in Supp. Table 2 were grown overnight on GCB-K. After 16-20 hours, strains were scraped into 0.3M sucrose, electroporated (V=1.8 kV, exponentially decaying wave) with 200 ng of plasmids DRE77-82, and rescued with GCP medium (15 g/L proteose peptone 3 (ThermoFisher), 1 g/L soluble starch (ThermoFisher), 4 g/L K2HPO4, 1 g/L KH2PO4, 5 g/L NaCl)^31^ with Kellogg’s supplement. After 10 minutes of rescue, transformants were then plated on nonselective GCB-K agar for 4 hours followed by selection on GCB-K supplemented with 70 μg/mL kanamycin. Finally, single colonies were re-streaked on GCB-K agar and checked for GyrA allele by Sanger sequencing. For GyrB cloning, fragments of GyrB were amplified using primers DR542 and DR543. Electroporation was performed as above, and single colonies were selected on GCB-K agar supplemented with 0.5 μg/mL ciprofloxacin.

### Genomic analysis

The genomic pipeline was conducted with methodology as published.^32^ Reads and metadata for the sequenced *N. gonorrhoeae* isolates were accessed from multiple sources^33–62^. Reads were then inspected using FastQC (v0.11.7) (https://www.bioinformatics.babraham.ac.uk/projects/fastqc/) and removed based on divergent GC content (from ~52-54%) or poor base quality. The remaining reads were mapped to the genome of *N. gonorrhoeae* strain NCCP11945 (RefSeq accession NC_011035.1) with BWA-MEM (v0.7.17-r1188)^63^. Mapped reads were then deduplicated with Picard (v2.20.1) (https://github.com/broadinstitute/picard). Samples with <40% coverage or <70% reads aligned were discovered with BamQC in Qualimap (v2.2.1)^64^ and discarded. Variants, including those in *gyrA* codons 91 and 95, were called with Pilon (v1.23)^65^ with a minimum mapping quality of 20 and a minimum depth of 10. Pseudogenomes were generated from the resulting Pilon VCFs with inclusion of all PASS sites and alternate alleles with allele frequency greater than 0.9. All other sites were set to “N”, and finally samples with >15% of sites called as missing were excluded. SPAdes (v3.12.0, with 8 threads, the –careful flag, and paired end reads where available)^66^ was used to create *de novo* assemblies, followed by contig filtering for quality (coverage >10X, length >500 bp, total genome size ~2.0-2.4 Mbp). These assemblies were annotated using Prokka (v1.14.6)^67^. Finally, the phylogenetic tree in this study was assembled in a recombination-corrected manner using Gubbins (v2.3.4)^68^ from all sequenced isolates from Reimche *et al*.^55^, representing lineages recently transmitting in the US, as well as the five genetically manipulated strains in Table 1 and all strains that had GyrA^91S^ and GyrA^95G/A/N^. The phylogeny was visualized in iTOL (v6)^69^.

**Table 1.**
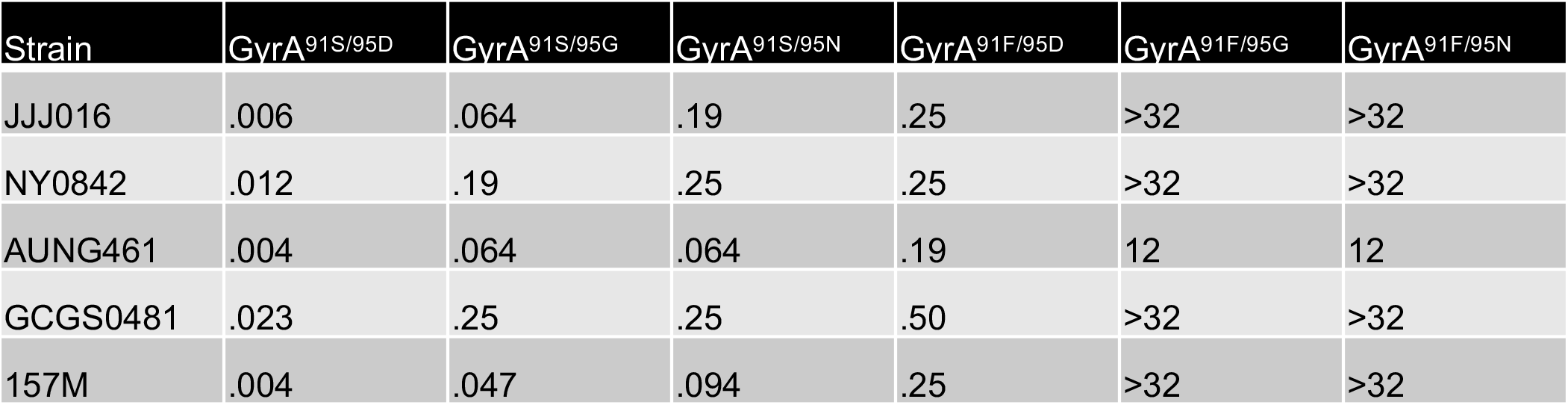
Ciprofloxacin MICs in μg/mL of five clinical isolates of *N. gonorrhoeae* following introduction of GyrA alleles via kanamycin co-selection.

| Strain | GyrA<sup>91S/95D</sup> | GyrA<sup>91S/95G</sup> | GyrA<sup>91S/95N</sup> | GyrA<sup>91F/95D</sup> | GyrA<sup>91F/95G</sup> | GyrA<sup>91F/95N</sup> |
| --- | --- | --- | --- | --- | --- | --- |
| JJJ016 | .006 | .064 | .19 | .25 | >32 | >32 |
| NY0842 | .012 | .19 | .25 | .25 | >32 | >32 |
| AUNG461 | .004 | .064 | .064 | .19 | 12 | 12 |
| GCGS0481 | .023 | .25 | .25 | .50 | >32 | >32 |
| 157M | .004 | .047 | .094 | .25 | >32 | >32 |

Transformants in this study were sequenced at SeqCenter (seqcenter.com). Sample libraries were assembled using the Illumina DNA Prep Kit and IDT 10bp UDI indices. The resulting libraries were sequenced with an Illumina NextSeq 2000. The paired-end 2×151 bp reads were demultiplexed, followed by quality control and adapter trimming, using bcl-convert (v3.9.3) (https://support.illumina.com/sequencing/sequencing_software/bcl-convert.html). Resulting reads were merged using BBMerge (v38.97)^70^. Reads were mapped to the parental genomic DNA assembly and variants predicted in Geneious 2021.0 (www.geneious.com) using default settings.

### Experimental evolution

Strain GCGS0481_SG (Supp. Table 1) was grown overnight on GCB-K agar. The overnight streak was scraped, diluted to ~OD 0.1, then plated as 14 replicates overnight onto GCB-K agar supplemented with 0.125 μg/mL ciprofloxacin. Each replicate was sequentially passaged, following the same dilution pattern, onto double the concentration of ciprofloxacin overnight until there was no visible growth or until 2 μg/mL was reached. Single colonies from the last resulting viable passage were selected for further analysis.

## Results

We selected five clinical *N. gonorrhoeae* isolates (Supp. Table 2) based on three major factors: high-level ciprofloxacin resistance, genetic and geographic diversity, and varying sets of resistance alleles (Figure 1A). All five of the strains encode GyrA^S91F^ as well as an additional mutation at GyrA position 95. All five strains carried mutations in ParC that are known to cause reduced susceptibility to ciprofloxacin, including ParC^S87R/N^ and ParC^E91G/K 71,72^. Finally, all five strains had the GyrB^429D^ allele associated with zoliflodacin susceptibility^26^.

**Figure 1.**
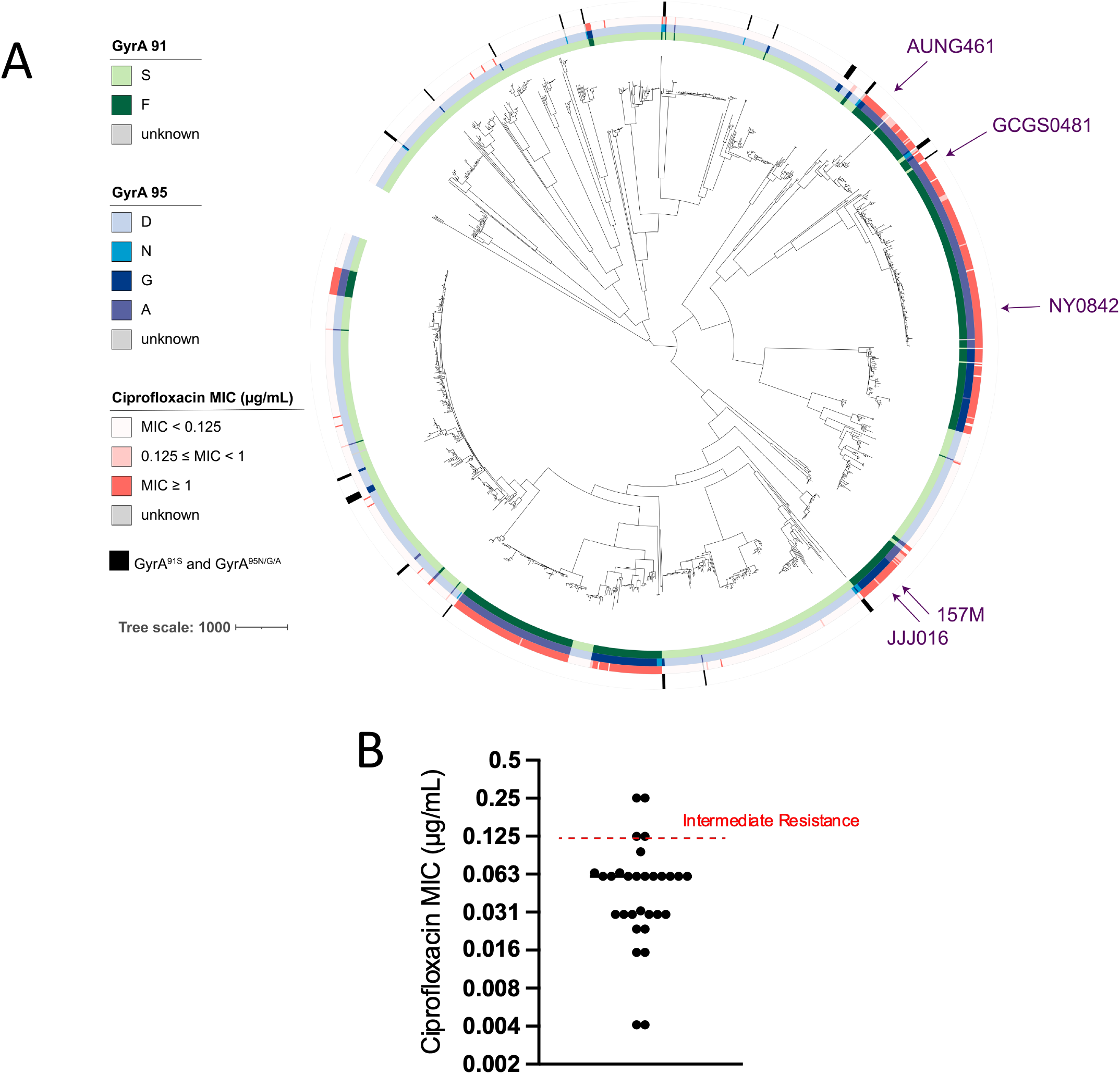
**(A)** Recombination-adjusted phylogenetic tree of 1,510 *N. gonorrhoeae* clinical isolates. On the innermost ring, the codon at position 91 of GyrA is coded by color, while on the second ring, the codon at position 95 of GyrA is similarly marked. The ciprofloxacin MIC of each isolate is shown on the third ring in red, while isolates with GyrA^91S^ and GyrA^85N/G/A^ are shown on the outermost ring in black. Isolates from this study are marked by arrows in purple. Tree scale represents recombination-adjusted number of SNPs. **(B)** Ciprofloxacin MICs of GyrA^91S^ and GyrA^95N/G/A^ strains indicated in **(A)**, with the CLSI threshold for intermediate resistance marked by a red dashed line.

We introduced six different alleles of GyrA into each strain using kanamycin co-selection, including pairwise combinations of codon 91 (91S – susceptible, 91F – resistant) and codon 95 (95D – susceptible, 95G, 95N – resistant). We were particularly interested in strains with the 91S codon, as these isolates would be reported as ciprofloxacin susceptible by assays focused on this codon. In all backgrounds, introduction of the GyrA^91S/95D^ allele, containing the susceptible variant at both codon 91 and 95, conferred susceptibility to ciprofloxacin. There was substantial MIC variation even at this baseline, with MICs varying almost 10-fold (Table 1).

Introduction of a mutation at the 95 codon, either GyrA^95G^ or GyrA^95N^, increased the MIC in all backgrounds by approximately 10-fold. Some isolates, including NY0842, a 2013 urethral isolate from New York City, and GCGS0481, a 2006 urethral isolate from U.S. Department of Health and Human Services Region 10 (Alaska, Idaho, Oregon, and Washington), reached an intermediate level of ciprofloxacin resistance by the Clinical and Laboratory Standards Institute (CLSI) breakpoint (≥0.125 μg/mL)^73^. Notably, MICs for ciprofloxacin ≥0.125 μg/mL are associated with substantially increased risk of treatment failure^74^. Finally, introduction of GyrA^91F/95G/N^ into all five backgrounds did not substantially affect MICs compared to the original isolates (Table 1, Supp. Table 2), indicating that the co-selected kanamycin cassette minimally affected the ciprofloxacin resistance profile.

These data suggest that some clinical isolates still retain ciprofloxacin non-susceptibility even after reversion of the GyrA^91F^ allele to GyrA^91S^. To evaluate whether examples with this genotype and phenotype have been observed, we investigated *gyrA* genotypes in 11,355 clinical isolates with published genomes and ciprofloxacin MICs. We identified 30 isolates bearing GyrA^91S/95G/N/A^, distributed across the *N. gonorrhoeae* phylogeny as singletons and closely related isolates (Figure 1A), suggesting that the GyrA^91S/95G/N/A^ alleles can arise in multiple genetic backgrounds and are sufficiently fit for transmission. The reported MICs for these isolates varied from 0.023 μg/mL to 0.25 μg/mL, including four with MICs above the clinically defined threshold where treatment failure is likely to occur (Figure 1B). Thus, diagnostic escape could occur either through reversion or expansion of specific clades with GyrA^91S^ and reduced susceptibility to ciprofloxacin.

We investigated the possibility that GyrA^91S^ isolates could develop higher level ciprofloxacin resistance through experimental evolution. We focused on the GyrA^91S/95G^ mutant of GCGS0481, differing from the parental isolate by only the kanamycin cassette and the 91S allele of *gyrA* as validated by whole-genome sequencing (WGS). We sequentially passaged GCGS0481 GyrA^91S/95G^ on increasing levels of ciprofloxacin, starting at ½ MIC (0.125 μg/mL) and increasing two-fold up to 2 μg/mL. Of the 14 parallel cultures, 10 (78%) had no viable colonies at the MIC. Single colonies from the four remaining cultures were then re-streaked and tested for ciprofloxacin susceptibility. One colony had a high ciprofloxacin MIC (>32 μg/mL), attributable to an S91F mutation; this colony was excluded from further analysis.

All three remaining mutants were isolated at 1 μg/mL ciprofloxacin, suggesting decreased susceptibility as compared to the GCGS0481 GyrA^91S/95G^ strain (0.25 μg/mL; Table 2) and clinical resistance to ciprofloxacin. WGS revealed that all three mutants had maintained GyrA^91S^ and acquired mutations in GyrB. One isolate had a missense mutation encoding a GyrB^E469D^ mutation. Introduction of this single-nucleotide polymorphism (SNP) in the parental isolate recapitulated the decreased susceptibility to ciprofloxacin and also conferred a two-fold increase in MIC for zoliflodacin (Table 2).

**Table 2.** Ciprofloxacin and zoliflodacin MICs of *N. gonorrhoeae* isolates with GyrA^91S/95G^ following experimental evolution in the presence of ciprofloxacin.

| Strain | Ciprofloxacin MIC (µg/mL) | Zoliflodacin MIC (µg/mL) | Ciprofloxacin concentration at isolation |
| --- | --- | --- | --- |
| GCGS0481 GyrA<sup>91F/95G</sup> (parent) | >32 | 0.25 | N/A |
| GCGS0481 GyrA<sup>91S/95G</sup> | 0.25 | 0.25 | N/A |
| GCGS0481 GyrA<sup>91S/95G</sup> Cip<sup>mut</sup> 1 | 1.5 | 0.5 | 1 µg/mL |
| GCGS0481 GyrA<sup>91S/95G</sup> GyrB<sup>E469D</sup> | 1.5 | 0.5 | N/A |
| GCGS0481 GyrA<sup>91S/95G</sup> Cip<sup>mut</sup> 2 | 1.5 | 2 | 1 µg/mL |
| GCGS0481 GyrA<sup>91S/95G</sup> Cip<sup>mut</sup> 3 | 1.5 | 2 | 1 µg/mL |
| GCGS0481 GyrA<sup>91S/95G</sup> GyrB<sup>D429N</sup> | 1.5 | 2 | N/A |

Two of the evolved strains gained GyrB^D429N^ mutations, which have been shown to cause decreased susceptibility to zoliflodacin^24,75^; we validated by WGS that these GyrB^D429N^ strains had not acquired other genomic mutations. Introduction of the GyrB^D429N^ mutation into the parental isolate recapitulated the phenotype of decreased susceptibility to ciprofloxacin and led to an 8-fold increase in the MIC for zoliflodacin as compared to the wildtype and parental GCGS0481 isolates (Table 2). These changes in antibiotic susceptibility are specific to ciprofloxacin and zoliflodacin (Supp. Table 3), suggesting that the differences in MIC are not due to general mechanisms of reduced drug susceptibility.

## Discussion

Our results indicate that GyrB^D429N^ is readily acquired by *N. gonorrhoeae* in the presence of ciprofloxacin and that GyrB^D429N^ increases ciprofloxacin resistance in at least some backgrounds. As such, this study raises both conceptual and practical implications for the introduction of assays using *gyrA* codon 91 to infer ciprofloxacin susceptibility. From a conceptual standpoint, this work provides an example of how escape from a diagnostic used to guide antibiotic therapy can drive the acquisition and possible spread of novel resistance determinants and how these pathways may have unintended consequences for resistance profiles to other antibiotics^23^. The presence of multiple clinical isolates with GyrA^91S^, together with resistance-associated mutations at GyrA^95^, strongly suggest that reversion of codon 91 is viable for pathogenesis. Our findings also indicate that a test that detects both position 91 and position 95 would be less likely to select for reversion to the susceptible allele, as all of the GyrA^91S/95D^ strains in this study remained highly susceptible to ciprofloxacin. Though the GyrB^E469^ mutation identified in this study has not been observed in clinical isolates of *N. gonorrhoeae*, the corresponding GyrB^E466D^ mutation in clinical isolates of *Salmonella enterica* serovar Typhimurium confers decreased susceptibility to ciprofloxacin^76^. This finding provides further evidence consistent with the hypothesis that diagnostic escape from a *gyrA* 91S assay may provide evolutionary pressure to drive novel resistance mechanisms.

The finding that GyrB^D429N^ reduces susceptibility to both ciprofloxacin and zoliflodacin was unexpected. Earlier work on zoliflodacin showed that, in ciprofloxacin-resistant clinical isolates from China, introduction of GyrB^D429N^ led to a decrease in ciprofloxacin susceptibility or had no effect on ciprofloxacin MIC^26^. Other studies have suggested that there is no cross-resistance between ciprofloxacin and zoliflodacin in recently sampled clinical isolates^77,78^, likely due to the finding that GyrB^D429N^ mutations have not been identified in sequences from ~13,000 clinical isolates^79^. In contrast, enzymology suggests that ciprofloxacin has modestly decreased potency for the stability of cleaved complexes with GyrB^D429N^ as compared to GyrB^429D^ (a CC50, a measure of the potency for stability of cleaved complexes, of 3 +/- 0.4 for the mutant vs. 1.9 +/- 0.7 for the WT)^75^. The cleaved complex is the adduct of enzyme and DNA formed at DNA nicks that are unable to be repaired in the presence of fluoroquinolones. As these cleaved complexes are thought to be correlated with, if not the mechanism of, fluoroquinolone-mediated cell death^80^, these data are consistent with zoliflodacin-ciprofloxacin cross-resistance. Further work is required to understand the genetic background dependence of phenotypic outcomes following introduction of GyrB^D429N^. One intriguing possibility is that GyrB^D429N^ is epistatic to GyrA^S91F^ with respect to ciprofloxacin resistance and is therefore less likely to arise in highly ciprofloxacin resistant backgrounds.

The findings in this work have implications for the introduction of zoliflodacin, pending a successful phase III trial. From the perspective of zoliflodacin resistance, testing for the GyrB^D429N^ allele should be prioritized in locations where ciprofloxacin is used, as well as in settings where assays targeting *gyrA* codon 91 inform treatment decisions. From the perspective of ciprofloxacin use, the emergence of zoliflodacin resistant isolates may also lead to an increase in ciprofloxacin MIC. GyrB^429^ should therefore be considered amongst the panel of mutations contributing to ciprofloxacin resistance and informing antibiotic selection for clinical decision-making.

## Supporting information

Rubin_GyrA_Supp

## Acknowledgements

We thank the members of the Grad Lab, particularly Sam Palace and the remainder of the GC subgroup, for their thoughts on and support for this project. We thank SeqCenter (seqcenter.com) for sequencing isolates.

