## Supplementary material for "*Neisseria gonorrhoeae* diagnostic escape from a *gyrA*-based test for ciprofloxacin susceptibility can lead to increased zoliflodacin resistance": Rubin_GyrA_Supp

Supplementary Tables 1-3

### Supp. Table 1

| Plasmid/Strain | Property(ies) | Citation |
|---|---|---|
| **Plasmids** | | |
| DRE77 | pUC19 with AphA3 $Kan^{R}$ cassette + homology to clone $GyrA^{91S/95D}$ | This Study |
| DRE78 | pUC19 with AphA3 $Kan^{R}$ cassette + homology to clone $GyrA^{91F/95D}$ | This Study |
| DRE79 | pUC19 with AphA3 $Kan^{R}$ cassette + homology to clone $GyrA^{91S/95N}$ | This Study |
| DRE80 | pUC19 with AphA3 $Kan^{R}$ cassette + homology to clone $GyrA^{91F/95G}$ | This Study |
| DRE81 | pUC19 with AphA3 $Kan^{R}$ cassette + homology to clone $GyrA^{91S/95N}$ | This Study |
| DRE82 | pUC19 with AphA3 $Kan^{R}$ cassette + homology to clone $GyrA^{91F/95N}$ | This Study |
| **Strains** | | |
| JJJ016_SD | Clinical *N. gonorrhoae* isolate JJJ016, $Kan^{R}$, $GyrA^{91S/95D}$ | This Study |
| JJJ016_FD | Clinical *N. gonorrhoae* isolate JJJ016, $Kan^{R}$, $GyrA^{91F/95D}$ | This Study |
| JJJ016_SN | Clinical *N. gonorrhoae* isolate JJJ016, $Kan^{R}$, $GyrA^{91S/95N}$ | This Study |
| JJJ016_FG | Clinical *N. gonorrhoae* isolate JJJ016, $Kan^{R}$, $GyrA^{91F/95G}$ | This Study |
| JJJ016_SN | Clinical *N. gonorrhoae* isolate JJJ016, $Kan^{R}$, $GyrA^{91S/95N}$ | This Study |
| JJJ016_FN | Clinical *N. gonorrhoae* isolate JJJ016, $Kan^{R}$, $GyrA^{91F/95N}$ | This Study |
| NY0842_SD | Clinical *N. gonorrhoae* isolate NY0842, $Kan^{R}$, $GyrA^{91S/95D}$ | This Study |
| NY0842_FD | Clinical *N. gonorrhoae* isolate NY0842, $Kan^{R}$, $GyrA^{91F/95D}$ | This Study |
| NY0842_SN | Clinical *N. gonorrhoae* isolate NY0842, $Kan^{R}$, $GyrA^{91S/95N}$ | This Study |
| NY0842_FG | Clinical *N. gonorrhoae* isolate NY0842, $Kan^{R}$, $GyrA^{91F/95G}$ | This Study |
| NY0842_SN | Clinical *N. gonorrhoae* isolate NY0842, $Kan^{R}$, $GyrA^{91S/95N}$ | This Study |
| NY0842_FN | Clinical *N. gonorrhoae* isolate NY0842, $Kan^{R}$, $GyrA^{91F/95N}$ | This Study |
| AUNG461_SD | Clinical *N. gonorrhoae* isolate AUNG461, $Kan^{R}$, $GyrA^{91S/95D}$ | This Study |
| AUNG461_FD | Clinical *N. gonorrhoae* isolate AUNG461, $Kan^{R}$, $GyrA^{91F/95D}$ | This Study |
| AUNG461_SN | Clinical *N. gonorrhoae* isolate AUNG461, $Kan^{R}$, $GyrA^{91S/95N}$ | This Study |
| AUNG461_FG | Clinical *N. gonorrhoae* isolate AUNG461, $Kan^{R}$, $GyrA^{91F/95G}$ | This Study |
| AUNG461_SN | Clinical *N. gonorrhoae* isolate AUNG461, $Kan^{R}$, $GyrA^{91S/95N}$ | This Study |
| AUNG461_FN | Clinical *N. gonorrhoae* isolate AUNG461, $Kan^{R}$, $GyrA^{91F/95N}$ | This Study |
| GCGS0481_SD | Clinical *N. gonorrhoae* isolate GCGS0481, $Kan^{R}$, $GyrA^{91S/95D}$ | This Study |
| GCGS0481_FD | Clinical *N. gonorrhoae* isolate GCGS0481, $Kan^{R}$, $GyrA^{91F/95D}$ | This Study |
| GCGS0481_SN | Clinical *N. gonorrhoae* isolate GCGS0481, $Kan^{R}$, $GyrA^{91S/95N}$ | This Study |
| GCGS0481_FG | Clinical *N. gonorrhoae* isolate GCGS0481, $Kan^{R}$, $GyrA^{91F/95G}$ | This Study |
| GCGS0481_SN | Clinical *N. gonorrhoae* isolate GCGS0481, $Kan^{R}$, $GyrA^{91S/95N}$ | This Study |
| GCGS0481_FN | Clinical *N. gonorrhoae* isolate GCGS0481, $Kan^{R}$, $GyrA^{91F/95N}$ | This Study |
| 157M_SD | Clinical *N. gonorrhoae* isolate 157M, $Kan^{R}$, $GyrA^{91S/95D}$ | This Study |
| 157M_FD | Clinical *N. gonorrhoae* isolate 157M, $Kan^{R}$, $GyrA^{91F/95D}$ | This Study |
| 157M_SN | Clinical *N. gonorrhoae* isolate 157M, $Kan^{R}$, $GyrA^{91S/95N}$ | This Study |
| 157M_FG | Clinical *N. gonorrhoae* isolate 157M, $Kan^{R}$, $GyrA^{91F/95G}$ | This Study |
| 157M_SN | Clinical *N. gonorrhoae* isolate 157M, $Kan^{R}$, $GyrA^{91S/95N}$ | This Study |
| 157M_FN | Clinical *N. gonorrhoae* isolate 157M, $Kan^{R}$, $GyrA^{91F/95N}$ | This Study |

| **Strains cont.** | | |
|---|---|---|
| GCGS0481_SG $Cip^{mut}$ 1 | GCGS0481_SG, $Kan^{R}$, isolated following sequential passage from ciprofloxacin 1 μg/mL passage, encodes $GyrB^{E469D}$ | This Study |
| GCGS0481_SG $GyrB^{E469D}$ | GCGS0481_SG, $Kan^{R}$, $GyrB^{E469D}$ | This Study |
| GCGS0481_SG $Cip^{mut}$ 2 | GCGS0481_SG, $Kan^{R}$, isolated following sequential passage from ciprofloxacin 1 μg/mL passage, encodes $GyrB^{D429N}$ | This Study |
| GCGS0481_SG $Cip^{mut}$ 3 | GCGS0481_SG, $Kan^{R}$, isolated following sequential passage from ciprofloxacin 1 μg/mL passage, encodes $GyrB^{D429N}$ | This Study |
| GCGS0481_SG $GyrB^{D429N}$ | GCGS0481_SG, $Kan^{R}$, $GyrB^{D429N}$ | This Study |

| **Primers** | **Sequence (Annealing)** | **Template** |
|---|---|---|
| DR_487_GyrAGib_1_F | TCCGTAGGTGAACCTGCGGGCGGCTGCTCGGG | NG gDNA |
| DR_477_GyrAGib_1_R | CGGCGGATCCCGCAGACCTTGTCAAAGCCGA | NG gDNA |
| DR_478_GyrAGib_2_F | CGGCTTTGACAAGGTCTGCGGGATCCGCCGTC | AphA3 $Kan^{R}$ cassette |
| DR_479_GyrAGib_2_R | AACATGATTTAAATAACGCGTCGACGCTTTTTA | AphA3 $Kan^{R}$ cassette |
| DR_480_GyrAGib_3_F | CGACGCGTTATTTAAATCATGTTGCGGGAAAGC | NG gDNA |
| DR_496_GyrAGib_3_R | GCTAGTTATTGCTCAGCGGGCGGCCTGTTTTATAGCCT | NG gDNA |
| DR_194_puc19_Gib_F | CCGCTGAGCAATAACTAGCGGATCCCCGGGTACCG | pUC19 |
| DR_195_puc19_Gib_R | CCGCAGGTTCACCTACGGATCTAGAGTCGACCTGCAGG | pUC19 |
| DR_542_gyrB_F | CAAGCAAACCGGAAAGTTCG | NG gDNA |
| DR_543_gyrB_R | GAAACCGCCGGCGAC | NG gDNA |

**Supplementary Table 1.** Strains, plasmids, and primers used in this study.

### Supp. Table 2

| Strain Name | Accession | Country | Anatomic Site | Year | Parental GyrA allele | Ciprofloxacin MIC |
|---|---|---|---|---|---|---|
| JJJ016 | SRR16683798 | Hong Kong | Urethra | 2015 | $GyrA^{91F/95N}$ | >32 |
| NY0842 | ERR2631880 | United States | Urethra | 2013 | $GyrA^{91F/95A}$ | >32 |
| AUNG461 | SRR8559420 | Australia | Urethra | 2017 | $GyrA^{91F/95G}$ | 12 |
| GCGS0481 | ERR855135 | United States | Urethra | 2006 | $GyrA^{91F/95G}$ | >32 |
| 157M | SRR16683705 | Vietnam | Urethra | 2019 | $GyrA^{91F/95N}$ | >32 |

**Supplementary Table 2.** Strain metadata for clinical *N. gonorrhoeae* isolates genetically manipulated in this study.

### Supp. Table 3

| Antibiotic MIC | GCGS0481 GyrA$^{91S/95G}$ | GCGS0481 GyrA$^{91S/95G}$ GyrB$^{D429N}$ |
|---|---|---|
| Ceftriaxone | 0.008 μg/mL | 0.008 μg/mL |
| Erythromycin | 4 μg/mL | 4 μg/mL |
| Benzylpenicillin | 0.75 μg/mL | 0.75 μg/mL |
| Tetracycline | 1 μg/mL | 1 μg/mL |

**Supplementary Table 3**. MICs for GCGS0481 GyrA$^{91S/95G}$ and GCGS0481 GyrA$^{91S/95G}$ GyrB$^{D429N}$.
